# Phylogenomics reveals extensive misidentification of fungal strains from the genus *Aspergillus*

**DOI:** 10.1101/2022.11.22.517304

**Authors:** Jacob L. Steenwyk, Charu Balamurugan, Huzefa A. Raja, Carla Gonçalves, Ningxiao Li, Frank Martin, Judith Berman, Nicholas H. Oberlies, John G. Gibbons, Gustavo H. Goldman, David M. Geiser, David S. Hibbett, Antonis Rokas

## Abstract

Modern taxonomic classification is often based on phylogenetic analyses of a few molecular markers, although single-gene studies are still common. However, the use of one or few molecular markers can lead to inaccurate inferences of species history and errors in classification. Here, we leverage genome-scale molecular phylogenetics (phylogenomics) of species and populations to reconstruct evolutionary relationships in a dense dataset of 711 fungal genomes from the biomedically and technologically important genus *Aspergillus*. To do so, we generated a novel set of 1,362 high-quality molecular markers specific for *Aspergillus* and provide profile Hidden Markov Models for each, facilitating others to use these molecular markers. Examination of the resulting genome-scale phylogeny: (1) helped resolve ongoing taxonomic controversies and identified new ones; (2) revealed extensive strain misidentification, underscoring the importance of population-level sampling in species classification; and (3) identified novel lineages that may shed light on the early evolution of an important genus. These findings suggest that phylogenomics of species and populations can facilitate accurate taxonomic classifications and reconstructions of the tree of life.

## Introduction

Accurate identification of fungal species underpins mycological research. Modern fungal taxonomy is based on diverse phenotypic traits and single or few molecular markers for species determination (Schoch et al., 2012; Houbraken et al., 2021). In addition, an increasing number of novel taxa, known from sequence data alone, are being discovered using environmental barcoding techniques (Lücking et al., 2021). For fungi, six molecular markers are used as universal barcodes: the ribosomal large and small subunit; ribosomal DNA array internal transcribed spacer region; the RNA polymerase II large subunit and core subunits; and minichromosome maintenance complex component 7 (Schoch et al., 2012; Raja et al., 2017). Two additional commonly used loci are transcription enhancer factor 1α and β-tubulin (Bruns et al., 1991; Hibbett et al., 2016; Raja et al., 2017; Lücking et al., 2021). Secondary barcodes are also often used to characterize species complexes (Bradshaw et al., 2022)—such as calmodulin for *Aspergillus* and other species (Susca et al., 2020)—owing to the power of more loci in phylogenetic reconstructions.

Typically, the sequences of one or more barcode loci are obtained from a strain of interest. Next, orthologous sequences are inferred based on sequence similarity from databases such as the National Center for Biotechnology Information; then, software is used to infer phylogenetic trees for each sequence, representing a hypothesis of the evolutionary history of these species (Raja et al., 2017). Despite the potential for genome-scale data to facilitate fungal taxonomy, current practices typically do not rely on whole-genome data, in part because of sparse genome sequences available across the fungal tree of life (Xu, 2020; Aime et al., 2021; Arias et al., 2021; Fountain et al., 2021; Kanamasa, 2021; Kück and Dahlmann, 2021; Li and Yu, 2021; Ruiz and Radwan, 2021; Stevenson and Stamps, 2021; Venkateswaran et al., 2021; Yin et al., 2021). Here, we tested the hypothesis that phylogenomic approaches that leverage whole genome sequences will improve the accuracy of phylogenetic reconstructions that have be based on sequences from a small number of loci (Rokas et al., 2003; Jarvis et al., 2014).

Fungi from the genus *Aspergillus* are of medical, agricultural, and biotechnological significance. *Aspergillus fumigatus* and *Aspergillus flavus* are pathogens, allergens, and produce mycotoxins (Nierman et al., 2005; Hedayati et al., 2007). *Aspergillus niger* is an industrial workhorse and *Aspergillus oryzae* is used in the production of fermented foods like soy sauce and sake (Gibbons et al., 2012; Cairns et al., 2018). Accurate identification of *Aspergillus* fungi with barcode loci is often challenging (Houbraken et al., 2021; Li and Yu, 2021; Stevenson and Stamps, 2021; Venkateswaran et al., 2021; Yin et al., 2021), but important because even closely related species can differ in drug resistance profiles and ability to cause disease (dos Santos et al., 2020; Steenwyk et al., 2020c). For example, clinical strains of *Aspergillus nomius* and *Aspergillus tamarii* have been misidentified as *Aspergillus flavus* (Tam et al., 2014). In the clinic, inaccurate species determination could lead to misguided disease management strategies due to differences in intrinsic drug resistance levels between species. For example, *A. latus*, commonly misidentified as *A. nidulans*, is more resistant to the antifungal drug caspofungin than *A. nidulans* (Steenwyk et al., 2020b).

Species are a taxonomic rank and serve as the foundation for comparative studies (De Queiroz, 2007; Houbraken et al., 2014; Xu, 2020; Aime et al., 2021). Several different species concepts have been proposed over the years (De Queiroz, 2007). Due to the microscopic nature of many fungi and lack of known sexual cycles for some species (Dyer and O’Gorman, 2012), species delimitation in the Kingdom Fungi has relied, in addition to cultural growth and micromorphological data, on molecular phylogenetics and the adoption of universal molecular barcodes (Schoch et al., 2012). Among *Aspergillus* fungi, multi-locus molecular phylogenetic approaches have become the predominant method (Samson et al., 2007, 2014; Houbraken and Samson, 2011; Houbraken et al., 2014). However, there is no current consensus for barcode similarity to designate a new species of fungi; for example, calmodulin gene sequences of *Aspergillus labruscus* and *Aspergillus oerlinghausenensis—*two recently described species of *Aspergillus—*share 85% and 97.3% sequence similarity to their closest relatives, *Aspergillus homomorphus* and *Aspergillus fumigatus*, respectively (Houbraken et al., 2016; Fungaro et al., 2017).

Reconstructing deeper evolutionary relationships from a few molecular markers can also be challenging. For example, divergences among sections—a secondary taxonomic rank above the species and below the genus ranks—have been debated. For example, the sections *Nigri, Ochraceorosei, Flavi, Circumdati, Candidi*, and *Terrei* were inferred to be monophyletic based on analyses of four loci from 81 taxa (Pitt and Taylor, 2014; Taylor et al., 2016) but topology tests using a 1,668-gene matrix from a different 81-taxon dataset rejected monophyly of these lineages (Steenwyk et al., 2019). Accurate reconstructions of the *Aspergillus* phylogeny will enable more accurate strain identification and will facilitate our understanding of how biomedically and technologically relevant traits—such as antimicrobial resistance—evolved.

Here, we present a dense phylogenomic tree inferred from a 1,362-gene matrix from 711 *Aspergillus* genome sequences spanning 106 species and population data for 36 species, more than doubling the number of species analyzed in previous genome-scale studies (de Vries et al., 2017; Steenwyk et al., 2019; Kjærbølling et al., 2020) and capturing roughly one quarter of all known species in the genus (Balajee, 2009; Sayers et al., 2009). The new phylogeny yields three key insights by: (1) demonstrating that population-level sequencing can facilitate strain classification; (2) resolving taxonomic controversies and identifying new ones; and (3) identifying a previously uncharacterized *Aspergillus* lineage that holds promise for elucidating the early evolution of this biomedically and technologically important genus.

## Results and Discussion

### A phylogenomic tree of *Aspergillus*

The evolutionary history of 725 genomes (711 *Aspergillus* genomes; 14 outgroup *Penicillium* genomes) was reconstructed using maximum likelihood analysis of a 1,362-gene data matrix with 6,378,237 nucleotide sites (Fig 1; Fig S1). The 725 genomes represent publicly available whole genome assemblies available through the National Center for Biotechnology Information (NCBI, https://www.ncbi.nlm.nih.gov/) that also passed quality-control measures (see *Methods*; Table S1 & S2). Based on the metadata provided to NCBI, the dataset includes 115 *Aspergillus* and 14 *Penicillium* species. The genomes of two or more strains were available for 36 *Aspergillus* species, but the depth of strain sampling varied (Table S3). Sampling was densest for *Aspergillus fumigatus* (N = 275), *Aspergillus flavus* (N = 105), *Aspergillus oryzae* (N = 97), and *Aspergillus niger* (N = 24). Sixteen species had genome sequences from two representative strains available. Twelve strains were of unknown species but were reported to be from the genus *Aspergillus*.

**Figure 1.**
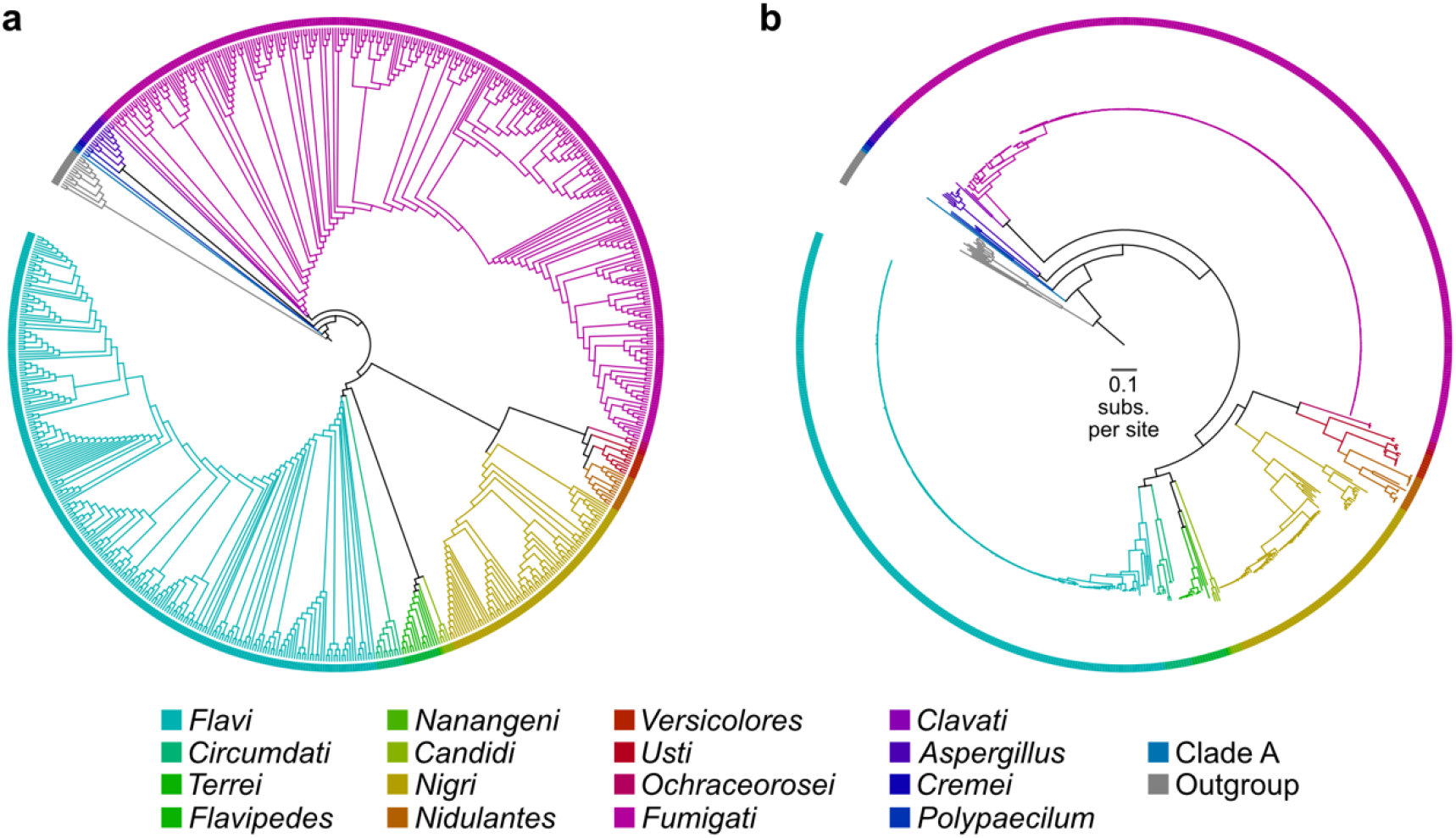
Phylogenomic tree of 725 genomes based on analysis of 1,362 genes (6,378,237 nucleotide sites). The evolutionary history of 711 *Aspergillus* species and 14 outgroup genomes from the genus *Penicillium* was reconstructed from a 1,362-gene matrix. The phylogeny is depicted without branch lengths (a) and with branch lengths, representing substitutions per site (b). Colors represent different sections—taxonomic ranks above species and below genus.

The dataset spanned 17 *Aspergillus* sections. Higher-order relationships among sections were examined using both concatenation- and coalescence-based tree inference approaches (Rokas et al., 2003; Mirarab et al., 2014). The phylogenies inferred with both approaches were highly congruent, differing by only one bipartition (Fig 2). Both approaches revealed the presence of a previously undescribed lineage, herein termed *Clade A* and represented by *Aspergillus* sp. MCCF 102 (Lekshmi et al., 2020). This lineage, by virtue of its placement as sister to all other sampled *Aspergillus* genomes, holds promise for understanding the early evolution of the genus *Aspergillus*.

**Figure 2.**
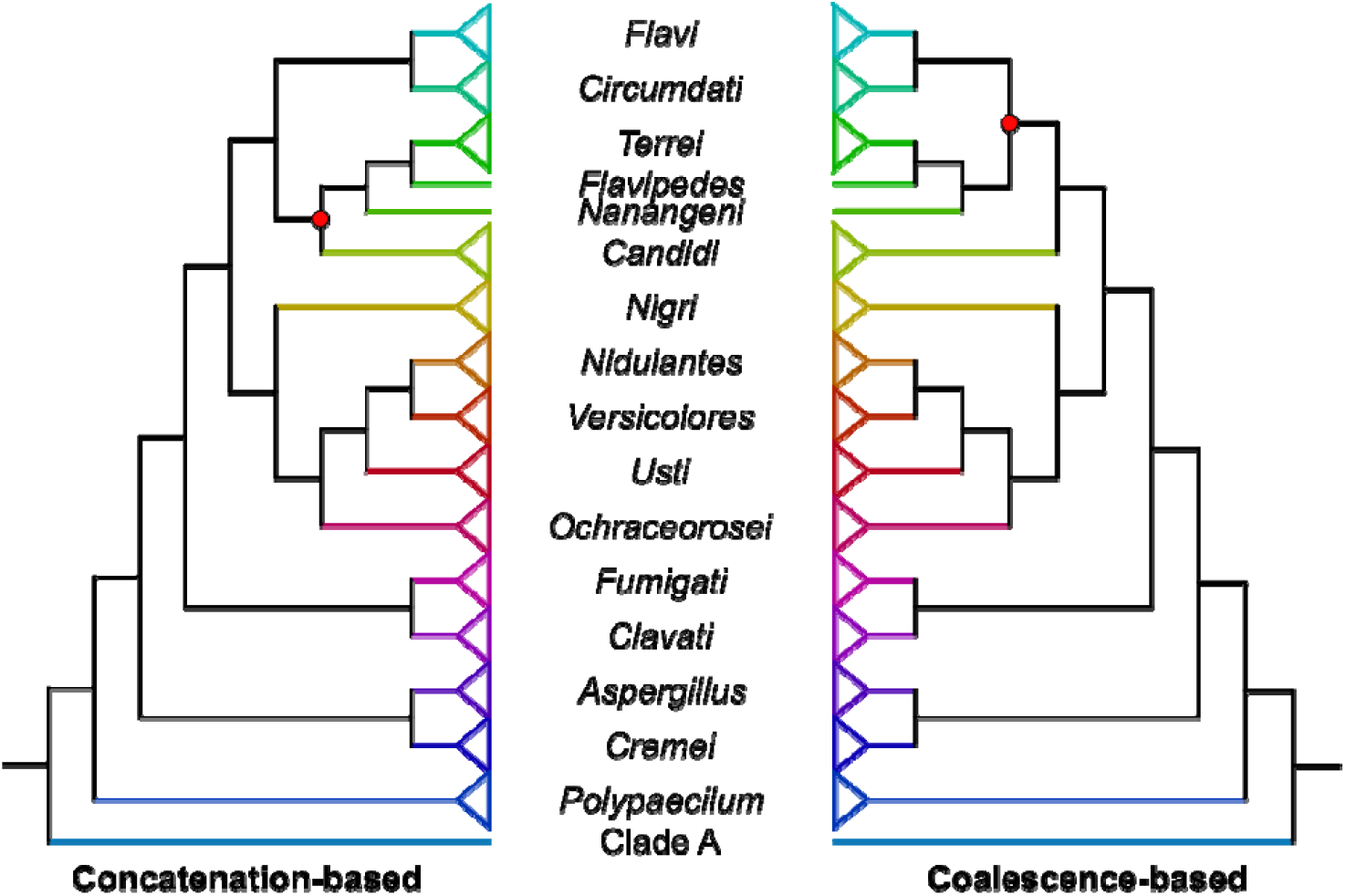
Concatenation- (left) and coalescence-based (right) phylogenies of taxonomic sections in the genus *Aspergillus* are highly congruent. The evolutionary relationships among sections are depicted. Species-level concatenation- and coalescence-based phylogenies differed at two bipartitions (represented by a red dot). Branch lengths and triangle sizes have no meaning.

### Phylogenomics sheds new light on *Aspergillus* strain identification

Phylogenomic analyses of the 711 strains reveal that half of the 36 species with multiple strains were not monophyletic (Table S4). Although monophyly is not required for species delimitation under certain species concepts (Rieseberg and Brouillet, 1994; Rieppel, 2010), this observation was surprising and suggests that cryptic speciation and / or species misidentification may be rampant. Here, we highlight six cases that illustrate this issue.

The phylogenetic placement of *Aspergillus neoellipticus* has been debated and bears on disease management strategies. Some studies suggest that *A. neoellipticus* is distinct from the major human pathogen *Aspergillus fumigatus*, based on the analysis of five loci (Li et al., 2014). Other studies suggested that *A. neoellipticus* is conspecific to *A. fumigatus* based on partial single-gene sequences and random amplified polymorphic DNA (Hong et al., 2005) (Peterson, 2008; Li et al., 2014, 2018). Phylogenomic analyses of species in section *Fumigati* (including four strains of *A. fumigatus* and one strain of *A. neoellipticus*) failed to resolve this ongoing debate, due to inferring *A. neoellipticus* as sister to *A. fumigatus* (Mead et al., 2021). The combined use of genome-scale data and extensive population-level sampling of 275 *A. fumigatus* strains in the analyses found that *A. neoellipticus* NRRL5109 is nested within a clade of 16 isolates of *A. fumigatus* (Fig 3a); the remaining 261 *A. fumigatus* isolates form the sister lineage.

**Figure 3.**
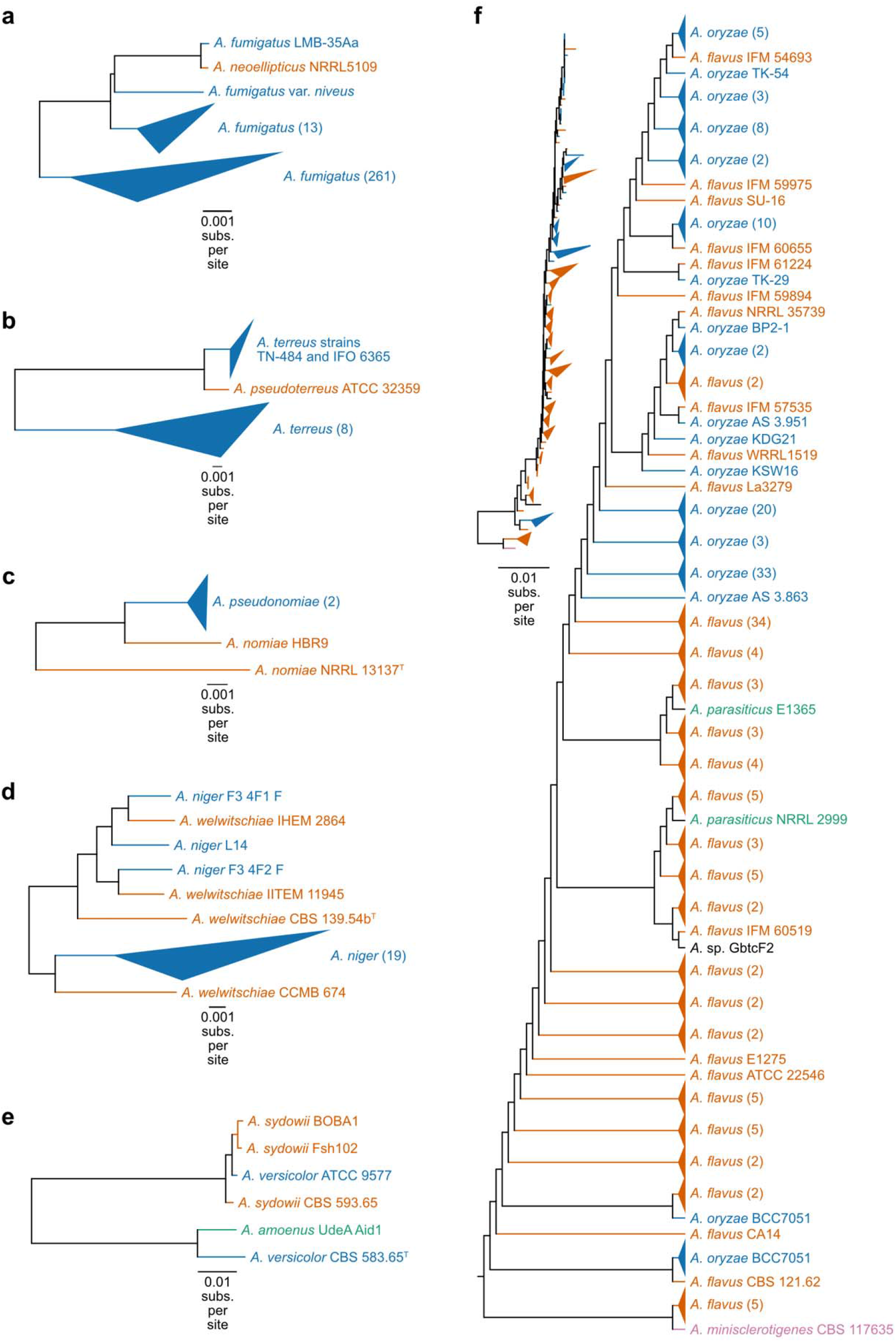
Phylogenomics underscores known taxonomic uncertainties and reveals new ones. (a) *Aspergillus neoellipticus* is a misidentified strain of *Aspergillus fumigatus*; alternatively, several *A. neoellipticus* strains are misidentified as *A. fumigatus*. (b) Strain misidentification occurs between *Aspergillus pseudoterreus* and *Aspergillus terreus* and (c) *Aspergillus pseudonomiae* and *Aspergillus nomiae*. (d) Isolates identified as *Aspergillus niger* and *Aspergillus welwitschiae* are likely misidentified. (e) *Aspergillus* strain ATCC 9577 is misidentified as *Aspergillus versicolor* but is likely *Aspergillus sydowii*. (f) Strains of *Aspergillus oryzae, Aspergillus flavus, Aspergillus parasiticus*, and *Aspergillus minisclerotigenes* appear to have polyphyletic origins, a result that is likely due to extensive strain misidentification (e.g., see also Houbraken et al 2021, detailing misidentification of five strains of *A. minisclerotigenes* as *A. flavus* (Houbraken et al., 2021)). Topologies presented were inferred using the concatenation approach. See Figure S3 for topologies inferred using coalescence. Different colors represent different species. Isolates with no known species are depicted in black. Triangles represent collapsed linages with multiple isolates. The number of isolates in each collapsed lineage are shown next to the species name in parentheses.

These results can be interpreted in two ways. Either *A. neoellipticus* NRRL5109 is an isolate of *A. fumigatus* or the clade formed by the 16 *A. fumigatus* strains and *A. neoellipticus* NRRL5109 is, in fact, *A. neoellipticus*, a species sister to *A. fumigatus*. If the latter, this would suggest that the 16 *A. fumigatus* isolates that form a clade with *A. neoellipticus* should be renamed to *A. neoellipticus*. Either way, these results underscore the importance of dense population-level representation for accurate strain identification.

A similar situation, but with much lower levels of sampling, exists for two other pairs of species: *Aspergillus terreus* and *Aspergillus pseudoterreus* (Fig 3b) and *A. nomiae* and *A. pseudonomiae* (Fig 3c). For *A. terreus* strains TN-484 and IFO 6365, our data corroborate recent findings that these are misidentified and should be *A. pseudoterreus*, suggesting misidentified information has persisted in databases (Houbraken et al., 2021).

Species from section *Nigri* are similar both in their morphology and in their molecular barcode sequences (Susca et al., 2016). *Aspergillus niger*, a species from the section *Nigri* (Coutinho et al., 2006), has been considered a prominent pathogen of sisal (*Agave sisalana*), an industrial crop used in the textile industry (Duarte et al., 2018). However, recent molecular phylogenetic analysis suggests that the main etiological agent of sisal bole rot disease is *Aspergillus welwitschiae*, rather than *A. niger* (Duarte et al., 2018). Examination of the evolutionary history of *A. niger* and *A. welwitschiae*, using phylogenomic analyses that sampled multiple strains from both species, identified four putative cases of strain misidentification (Fig 3d). Three strains (F3 4F1 F, L14, and F3 4F2 F), previously identified as *A. niger*, are conspecific with the majority of *A. welwitschiae* strains, suggesting that they are, indeed, *A. welwitschiae* and not *A. niger*. The fourth strain, CCMB 674, which was previously identified as *A. welwitschiae*, is sister to a lineage of 19 *A. niger* strains; therefore, isolate CCMB 674 is likely *A. niger* and not *A. welwitschiae*. These findings suggest that strain misidentification confounds studies of sisal bole rot, which could have major economic ramifications.

Another instance of putative misidentification concerns *Aspergillus versicolor* ATCC 9577, which is conspecific with *Aspergillus sydowii* strains, suggesting that strain ATCC 9577 is in fact *A. sydowii*. In contrast, the type strain of *A. sydowii*, CBS 583.65^T^, is closely related to *A. amoenus*, an observation that is corroborated by previous taxonomic studies (Jurjevic et al., 2012). These findings suggest that multiple strains from multiple species may be misidentified. Furthermore, ATCC 9577 and CBS 583.65 are reportedly synonymous strain names (Sklenář et al., 2022); the fact that there are two different genome sequences available is suggestive of contamination or erroneous metadata for at least one of these strains. These findings suggest that the entire clade may benefit from taxonomic revision and closer scrutiny of strain identity. To this point, a recent analysis of a five-gene, 213-taxon dataset proposed species in the series *Versicolores* be reduced from 17 species to four, citing intraspecific variation as a driver of over splitting of species boundaries by taxonomists (Sklenář et al., 2022). Evolutionary relationships under this new analysis differ slightly from our genome-scale phylogeny (Fig S2). Additional genome sequences of species and strains may shed light on the evolutionary history of section *Versicolores*.

Accurate classification of *A. flavus, A. oryzae*, and *Aspergillus parasiticus* is important because of biomedical and food safety concerns (Fig 3f). *Aspergillus flavus* is a human pathogen, post-harvest food pathogen, and mycotoxin producer (Gourama and Bullerman, 1995; Geiser et al., 1998; Hedayati et al., 2007). *Aspergillus oryzae* is the domesticate of *A. flavus* used for sake production (Gibbons et al., 2012). Successive short branch lengths and non-monophyly of *A. flavus* and *A. oryzae* strains may suggest multiple domestication events or the introduction of domesticated isolates into the wild. Another interpretation is that the two species are distinct ecotypes (or populations), rather than distinct species. Notably, *A. oryzae* strains are known to produce fewer mycotoxins than *A. flavus* (Gibbons et al., 2012), which may be a diagnostic signature (i.e., phenotype) of an ecotype appropriate for food production. Interestingly, the aflatoxigenic postharvest pathogen *A. parasiticus* is conspecific with *A. flavus* strains, and the two *A. parasiticus* strains do not form a monophyletic group; strain NRRL 2999 is misidentified as *A. parasiticus* but is, in fact, *A. flavus* (Houbraken et al., 2021). Five isolates of *A. flavus* are sister to *A. minisclerotigenes* and more distantly related to other *A. flavus* isolates; our genome-scale analyses are consistent with inferences based on a recent examination of individual barcode loci (Houbraken et al., 2021) in suggesting that the five *A. flavus* isolates are misidentified. Lastly, five isolates with unknown species designations were determined, whereas six remain unknown (Table S5).

### A roadmap for studies of *Aspergillus*

We present a comprehensive genome-scale phylogeny of *Aspergillus* (Fig 1 and 4). Our results underscore the need for further research into *Aspergillus* species delimitation and that strain misidentification may be a more common problem among publicly available data than previously appreciated (Houbraken et al., 2021). Strain misidentification can be reduced by two factors: (1) genome-scale data, which are less prone to errors in phylogenetic inference compared to single or a few loci (Rokas et al., 2003; Jarvis et al., 2014; Steenwyk et al., 2019) and (2) population-level sampling from diverse environmental niches and geographic locations. Combined, these two factors facilitated identifying cases where strains may represent new species or should be conspecific with another (such as *A. neoellipticus*) or uncovered instances of more rampant strain misidentification (such as the case of *A. niger* and *A. welwitschiae* and species in section *Flavi*). Lastly, our analyses identified a new lineage (*Aspergillus* sp. MCCF 102) that is sister to all other *Aspergillus* strains whose genomes have been sequenced and may hold clues to the early evolution of *Aspergillus* species. To facilitate others using our findings, we summarize higher-order and species relationships among *Aspergillus* species (Fig 4).

**Figure 4.**
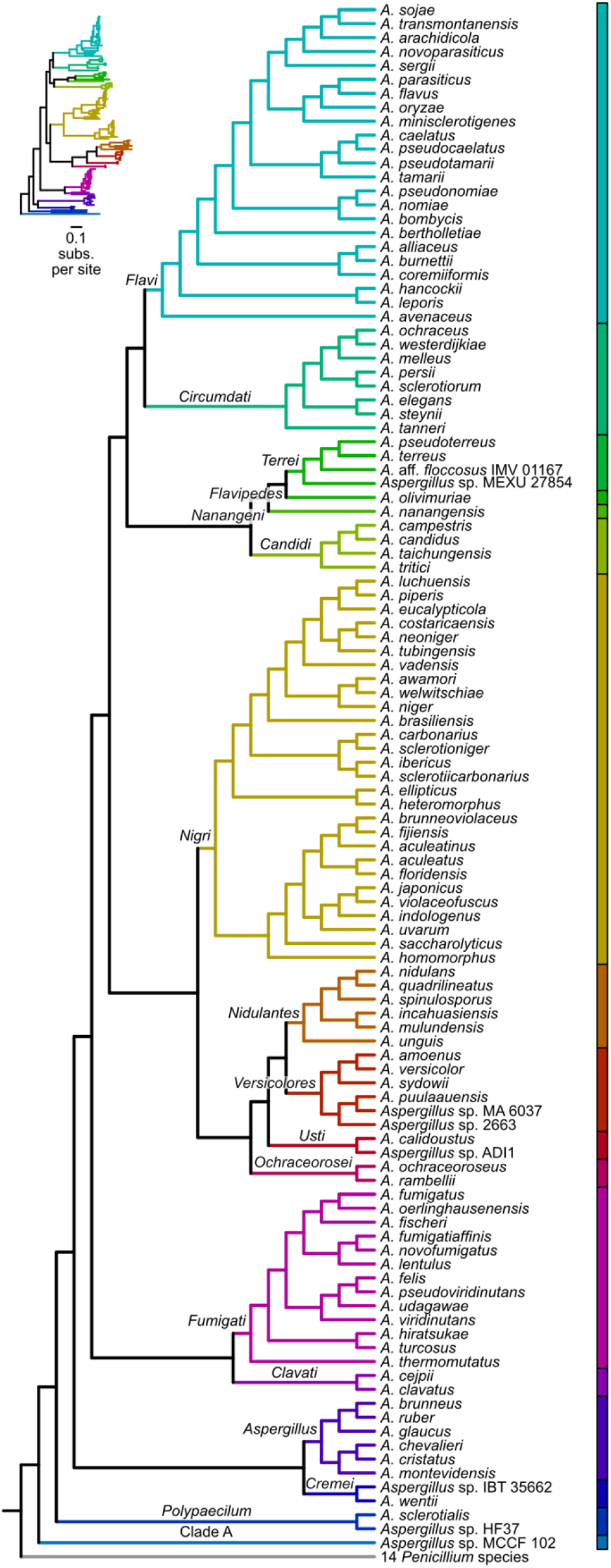
A species-level phylogeny of the genus *Aspergillus*. A genome-scale view of the *Aspergillus* phylogeny and the identification of a new sister lineage, clade A, to a clade of the rest of *Aspergillus* species whose genomes have been sequenced may help understand the early evolution of this biomedically and technologically relevant lineage. Inset represents the same phylogeny with branch lengths representing substitutions per site.

Resolving these issues and facilitating future identification of strains and species requires novel strategies and prioritizes analysis of type strains. Similar to databases of internal transcribed spacer regions of fungi that help facilitate isolate and strain classification (Nilsson et al., 2019), genome-scale resources can be utilized for strain classification, an approach that is of growing interest among budding yeast biologists (Libkind et al., 2020), but yet to be well adopted for other fungal lineages. To facilitate strain identification among *Aspergillus* species, we have produced profile Hidden Markov Models of the 1,362 molecular markers used in the present study (see *Data Availability*). Adding newly sequenced strains to our proposed phylogenetic tree of *Aspergillus* may be a helpful step toward accurate strain identification. Furthermore, additional research is needed to identify and characterize cryptic species, that is organisms that are morphologically highly similar to known species but genetically and physiologically distinct (Geiser et al., 1998; Alastruey-Izquierdo et al., 2014). Accurate strain identification and elucidation of species boundaries will greatly benefit from increased genome sequencing of type strains.

## Methods

### Genome data acquisition and quality control

All publicly available *Aspergillus* genomes (N = 717) were downloaded from NCBI (National Center for Biotechnology Information; https://www.ncbi.nlm.nih.gov/; date accessed: January 9^th^, 2022). Publicly available genome annotations were also downloaded. For genomes without available annotations, gene boundaries were predicted using AUGUSTUS, v3.3.2 (Stanke and Waack, 2003), with the “species” parameter set to “aspergillus_nidulans.” To determine if the genomes were suitable to phylogenomic analyses, gene prediction completeness was examined using BUSCO, v4.0.4 (Waterhouse et al., 2018), and the Eurotiales database of 4,191 near-universally single-copy orthologs (or BUSCO genes) from OrthoDB, v10 (Kriventseva et al., 2019). Six genomes with less than 75% single-copy complete BUSCO genes were removed, resulting in 711 *Aspergillus* genomes. The resulting sets of gene predictions were highly complete (mean ± standard deviation: 95.74 ± 2.25%). For outgroup taxa, 14 *Penicillium* genomes and annotations were downloaded from NCBI. The completeness of *Penicillium* gene predictions were assessed using the same protocol and were highly complete (mean ± standard deviation: 94.73 ± 4.04%). The final dataset had 725 genomes.

### Single-copy orthologous gene identification

Phylogenomics often relies on single-copy orthologous genes. OrthoFinder, v.2.3.8 (Emms and Kelly, 2019), was used to identify single-copy orthologous genes by clustering protein sequences into groups of orthologs. Clustering sequences was based on protein sequence similarity and calculated using DIAMOND, v2.0.13.151 (Buchfink et al., 2021). To reduce computation time and memory, orthology predictions were conducted among 40 representative species that span the diversity of *Aspergillus* species (Table S6) (Steenwyk et al., 2019). The impact of 41 different inflation parameter settings (one through five with a step of 0.1) on the number of single-copy orthologs identified was examined. The inflation parameter that resulted in the highest number of single-copy orthologs was 3.6. The resulting 7,882 single-copy orthologs with at least 50% occupancy (N = 20) were used for downstream analysis.

To identify orthologs in the full 725-genome dataset, sequence similarity searches were conducted in each proteome. To do so, the 7,882 single-copy orthologs were aligned using MAFFT, v7.402 (Katoh and Standley, 2013), with the auto parameter. Profile Hidden Markov Models (HMMs) were then built for each alignment using the hmmbuild function in HMMER, v3.1b2 (Eddy, 2011). The resulting HMMs were used to identify single-copy orthologs in the 725 proteomes using orthofisher, v.1.0.3 (Steenwyk and Rokas, 2021), and a bitscore fraction threshold of 0.95.

To generate single-gene phylogenies, the protein sequences of the 7,882 single-copy orthologs, identified using orthofisher (Steenwyk and Rokas, 2021), were aligned using MAFFT as described above (Katoh and Standley, 2013). The corresponding nucleotide sequences were threaded onto the protein alignment using the thread_dna function in PhyKIT, v1.11.12 (Steenwyk et al., 2021) and trimmed using ClipKIT, v1.3.0 (Steenwyk et al., 2020a). Excessive trimming of multiple sequence alignments worsens single-gene phylogenetic inference (Tan et al., 2015; Steenwyk et al., 2020a); thus, multiple sequence alignments wherein 40% or more of the original alignment length was maintained after trimming were retained resulting in 4,300 single-copy orthologs. The evolutionary histories of the 4,300 single-copy orthologs were inferred using IQ-TREE 2 (Minh et al., 2020, 2). The best-fitting substitution model was selected using ModelFinder (Kalyaanamoorthy et al., 2017). To remove potential instances of hidden paralogy, the monophyly of the five well-established lineages was examined using PhyKIT (Steenwyk et al., 2021). Specifically, the single-gene phylogenies were examined for the monophyly of five well established lineages: sections *Flavi* (N = 246), *Fumigati* (N = 316), *Nidulantes* (N = 12), and *Versicolores* (N = 7) as well as the outgroup lineage of 14 *Penicillium* species. Genes wherein one or more of the five lineages were not monophyletic were removed, resulting in a final set of 1,362 single-copy orthologous genes. The average occupancy for each single-copy ortholog was 0.98 ± 0.07.

### Phylogenomic tree inference

The final set of single-copy orthologs was used to infer the evolutionary history of the 725 species and strains. Specifically, the 1,362 single-copy orthologs were concatenated into a single matrix using the create_concat function in PhyKIT (Steenwyk et al., 2021). The resulting supermatrix had 6,378,237 sites (2,846,432 parsimony informative sites). Alignment length and number of parsimony informative sites were calculated using BioKIT, v0.1.2 (Steenwyk et al., 2022). IQ-TREE 2 was used for tree inference (Minh et al., 2020, 2). The best fitting substitution model was selected using ModelFinder (Kalyaanamoorthy et al., 2017). Bipartition support was assessed using 1,000 ultrafast bootstrap approximations (Hoang et al., 2018). The total central processing unit (or CPU) hours required for this analysis was 2,763 (approximately 4 months). An additional coalescence-based consensus tree approach was used to infer the evolutionary history among species and strains. Specifically, the 1,362 single-copy orthologous gene phylogenies were used as input to ASTRAL, v5.6.3 (Zhang et al., 2018). Phylogenies were visualized using iTOL, v6 (Letunic and Bork, 2019).

## Data availability

Results and data presented in this study will be available from figshare upon manuscript acceptance (doi: 10.6084/m9.figshare.21382131). The following link is provided for review purposes only https://figshare.com/s/6b91c4f3f575c0ef9883.

## Acknowledgements

We thank the Rokas lab for helpful discussion and comments.

