## Supplementary Figures for "Phylogenomics reveals extensive misidentification of fungal strains from the genus *Aspergillus*"

**Supplementary figures and supplementary figure legends**

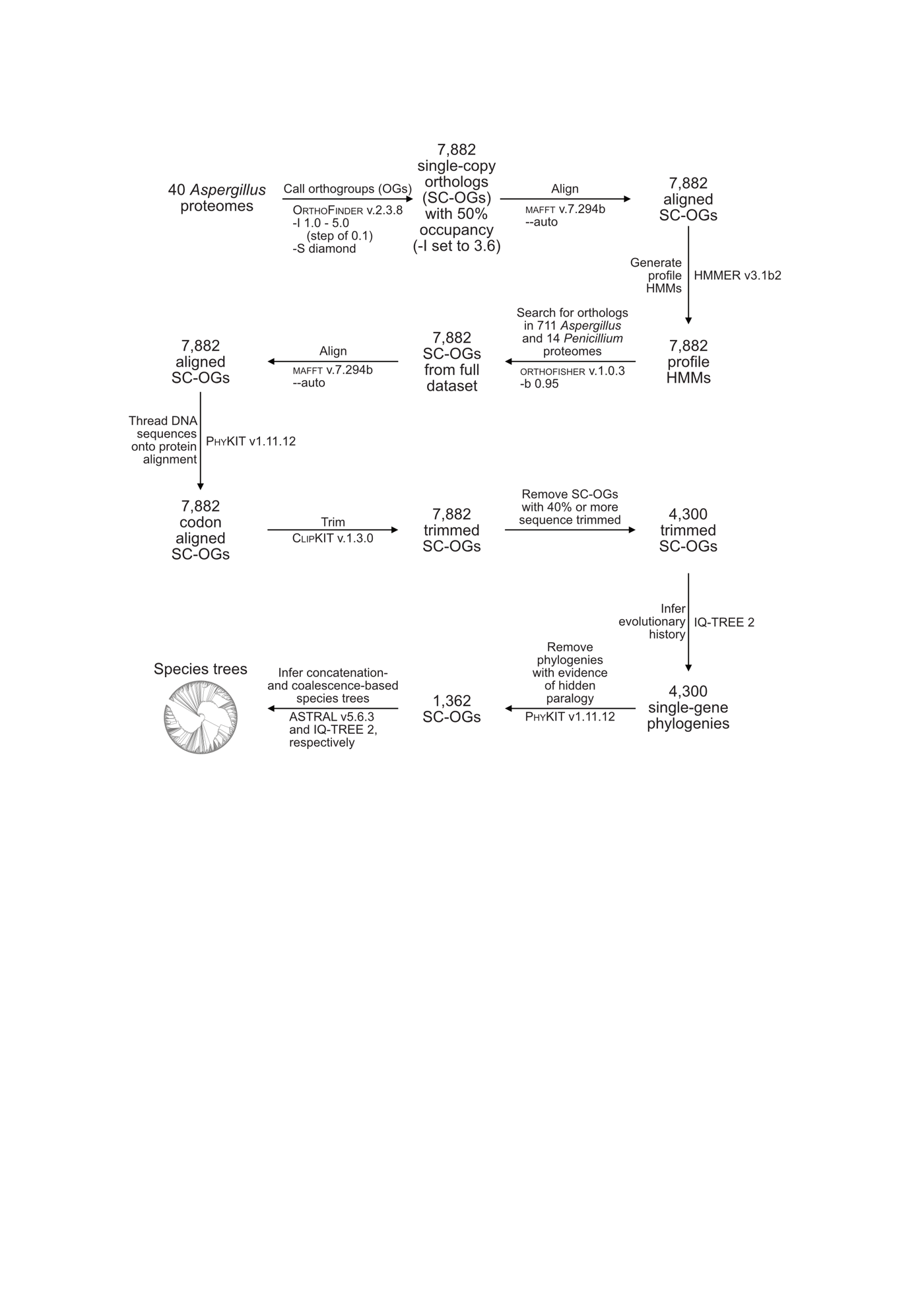

**Figure S1. Diagram of methods.** A diagrammatic representation of the methods is depicted here. Overall, the workflow represents an approach to identifying high-quality single-copy orthologous genes that serve as molecular markers for phylogenomic analysis. Key cleaning producers include removing poor quality markers due to insufficient taxon occupancy and evidence of hidden paralogy. This process resulted in 1,362 single-copy orthologous genes that were used as phylogenomic markers.

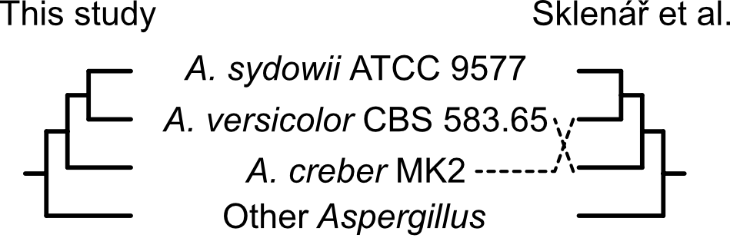

**Figure S2. Comparing relationships among species in section *Versicolores* under a new taxonomic schema proposed by Skelář *et al*. 2022.** The number of species in series *Versicolores* has recently been reduced from 17 to four (Sklenář et al., 2022), three of which are represented in the present study. Evolutionary relationships differ between the five-locus study and our genome-scale study. Although our study benefits from additional loci, which can help elucidate evolutionary relationships, our study also suffers from reduced taxon sampling (7 strains vs. 213). Moreover, our analysis did not include the type strain of *Aspergillus creber* due to a lack of genome sequence availability.

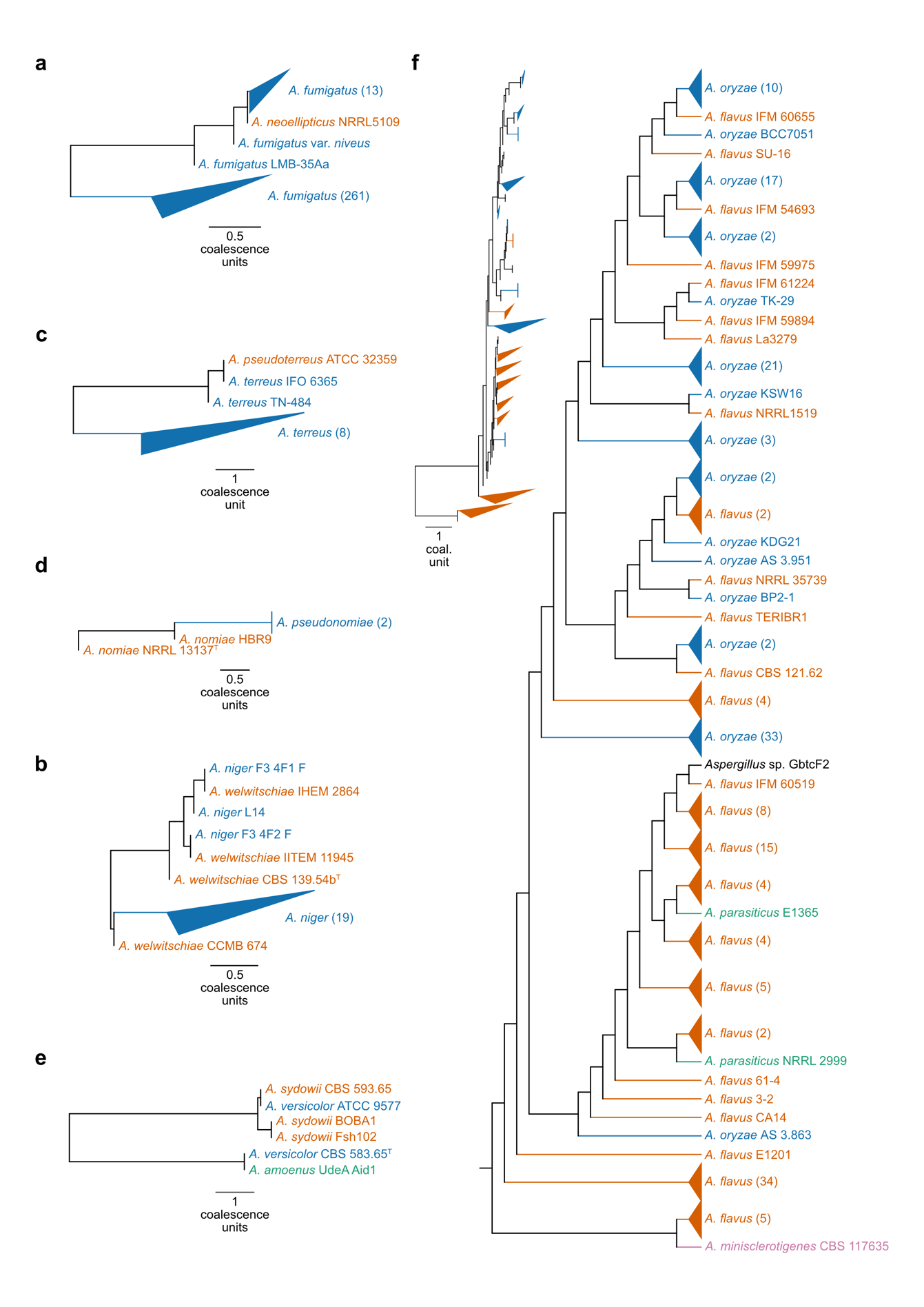

**Figure S3. Phylogenomics using coalescence underscores known taxonomic uncertainties and reveals new ones.** (a) *Aspergillus neoellipticus* is a strain of *Aspergillus fumigatus*; alternatively, several *A. neoellipticus* strains are misidentified as *A. fumigatus*. (b) Isolates identified as *Aspergillus niger* and *Aspergillus welwitschiae* are misidentified. (c) Strain misidentification occurs between *Aspergillus pseudoterreus* and *Aspergillus terreus* and (d) *Aspergillus pseudonomiae* and *Aspergillus nomiae*. (e) *Aspergillus* strain ATCC 9577 is misidentified as *Aspergillus versicolor* but is likely *Aspergillus sydowii*. (f) Strains of *Aspergillus oryzae*, *Aspergillus flavus*, *Aspergillus parasiticus*, and *Aspergillus minisclerotigenes* appear polyphyletic, a result that is likely due to extensive strain misidentification (see also Houbraken et al 2021 – (Houbraken et al., 2021)). Topologies presented were inferred using the concatenation approach. See Figure 3 for topologies inferred using concatenation. Different colors represent different species. Isolates with no known species are depicted in black. Triangles represent collapsed linages with multiple isolates. The number of isolates in each collapsed lineage are shown next to the species name in parentheses.
